# Altered neurogenic pathways in experimental rat models of ischemic cerebral stroke: A transcriptome based meta-analysis

**DOI:** 10.1101/2021.11.24.469795

**Authors:** Syed Aasish Roshan, Gayathri Elangovan, Dharani Gunaseelan, Swaminathan K. Jayachandran, Mahesh Kandasamy, Muthuswamy Anusuyadevi

**Author notes:** **Address for Correspondence:** Dr. Muthuswamy Anusuyadevi, Ph.D, Associate Professor, Department of Biochemistry, Molecular Neuro-Gerontology Laboratory, School of Life Sciences, Bharathidasan University, Tiruchirappalli – 620024, Tamilnadu, India.

## Abstract

1)

**Objectives:** Promoting neurogenesis mediated recovery is one of the most sought after strategies in recovery after cerebral stroke. In this paper we elucidate how neurogenesis related genes are altered in the early stroke environment, to hint at potential pathways for therapeutic recovery.

**Materials and Methods:** Around 97 microarray datasets derived from stroke affected rat brains were collected from NCBI-GEO. Datasets were normalized and subjected to meta-analysis in Network Analyst to identify differentially expressed genes. Gene enrichment analyses were carried out using GSEA, and WebGestalt, and results were visualized using Cytoscape Enrichment mapping.

**Results:** Nearly 939 differentially expressing genes were identified in the cerebral stroke group. Among them 30 neurogenesis related genes were identified through enrichment mapping analysis, and 35 genes through Protein- Protein Interaction analysis. Highest upregulated neurogenesis genes were found to be TSPO, GFAP, VIM, and TGFB1. Highest Downregulated neurogenesis genes were found to be THY1, NR1D1, CDK5, STX1B, and NOG.

**Conclusions:** Through this study, we have identified that during the acute time frame after stroke, majority of the neurogenesis genes related to neural proliferation and neural differentiation are downregulated, while the majority of the genes related to neuronal migration were upregulated. Single or combined therapeutic approach against the identified dysregulated genes could greatly aid neural restoration and functional recovery during the postischemic stage.

## 2) Introduction

Ischemic stroke is caused by an abrupt disruption in the blood flow and impairments in vascular compartments of the brain. The onset of ischemic stroke leads to the deprivation of oxygen and glucose in brain cells, especially neurons, as they have very poor energy reserves. The pathophysiological events following this blockage ultimately leads to neuronal loss in the infarct regions leaving the peri-infarct region vulnerable to necrotic and apoptotic cell death followed by fibrotic formations [1, 2]. As the majority of strokes occur due to abnormalities of the middle cerebral artery, the striatum appears to be highly vulnerable to ischemia among other brain regions. Hence, the prominent neuronal loss in the ischemic striatum leaves the patient with a major motor deficit or to a paralysed state [3]. Recombinant tissue plasminogen activator (rTPA) has been the effective management therapy available for recovery after ischemic stroke, however with a therapeutic window of fewer than 4.5 hours from the onset. This leaves no actual solution as a treatment option for stroke beyond the golden hours. Hence, establishment of valid treatment against MCAo stroke has been an important area in preclinical research, and it needs ample research to establish more appropriate therapeutic targets for the fruitful outcome.

Adult Neurogenesis is the innate ability of the brain to continuously produce new neuronal cells even till a later stage of life. Adult Neurogenesis occurs predominantly in two major stem cell niches of the brain, namely subgranular zone ( SGZ ) in the dentate gyrus of the hippocampus, and the subventricular zone ( SVZ ) present along the lateral ventricles of the brain. Experimental paradigms targeting the regulation of neurogenesis to promote neural regeneration as a treatment option against stroke is an ongoing scientific task for the past two decades. Despite the accumulation of reports, regenerative strategies against stroke are very feeble due to the lack of complete understanding in the critical aspects of signalling mechanisms.

With the recent scientific advancement in omics approaches, powerful tools like microarray analysis, RNA-seq and other Seq techniques have become easily accessible to the research community. The data from these experiments, though different in study design, but enables cross-comparison, provided appropriate control measures were taken. Though there is a huge accumulation of MCAo stroke gene expression data, the experimental designs in each study tend to focus on only a few specific scientific problems, it is still challenging to extract the maximal information on brain regeneration from those data. To address these issues, the establishment of a meta-analysis that integrates and inter-validates quantitative findings of multiple independent datasets from publicly available microarray data in databases including the Gene Expression Omnibus (GEO, HTTP:// ncbi.nlm.nih.gov/geo) would be highly instrumental, as it can offer increase statistical accuracy due to augmented sample size, control quality for between-study variation, and overcoming the bias of individual studies. The results from such an integrated meta-analysis experiment could help in further identification of potential biomarkers and druggable targets which could be further translated into diagnostic and treatment applications. Hence, this meta-analysis attempts to establish a potential list of differentially expressed genes, which could be further analysed to identify dysregulated signalling pathways that determine the regulation of adult neurogenesis from the available microarray data from rat stroke brain.

## 3) Materials and Methods

### 3.1) Collection of microarray data

A computerised search was carried out in NCBI GEO against the terms “MCAO” and “stroke” with the organism set as Rattus norwegicus. The microarray datasets of experimental cerebral stroke, and studies with similar experimental conditions were shortlisted. Inclusion criteria considered for the study have been supplied in the table-1. Details about the collected datasets are listed in table-2. Here, 97 samples (54 MCAo brain samples vs 43 sham control brain samples) were selected from 10 microarray datasets. An overview of the meta-analysis conducted in this study is shown in Figure-1. The studies collected utilized a range of microarray platforms like Agilent, Affymetrix, and Illumina. Throughout the study, only the ipsilateral cortex of the rat brain was considered.

**Figure-1:**
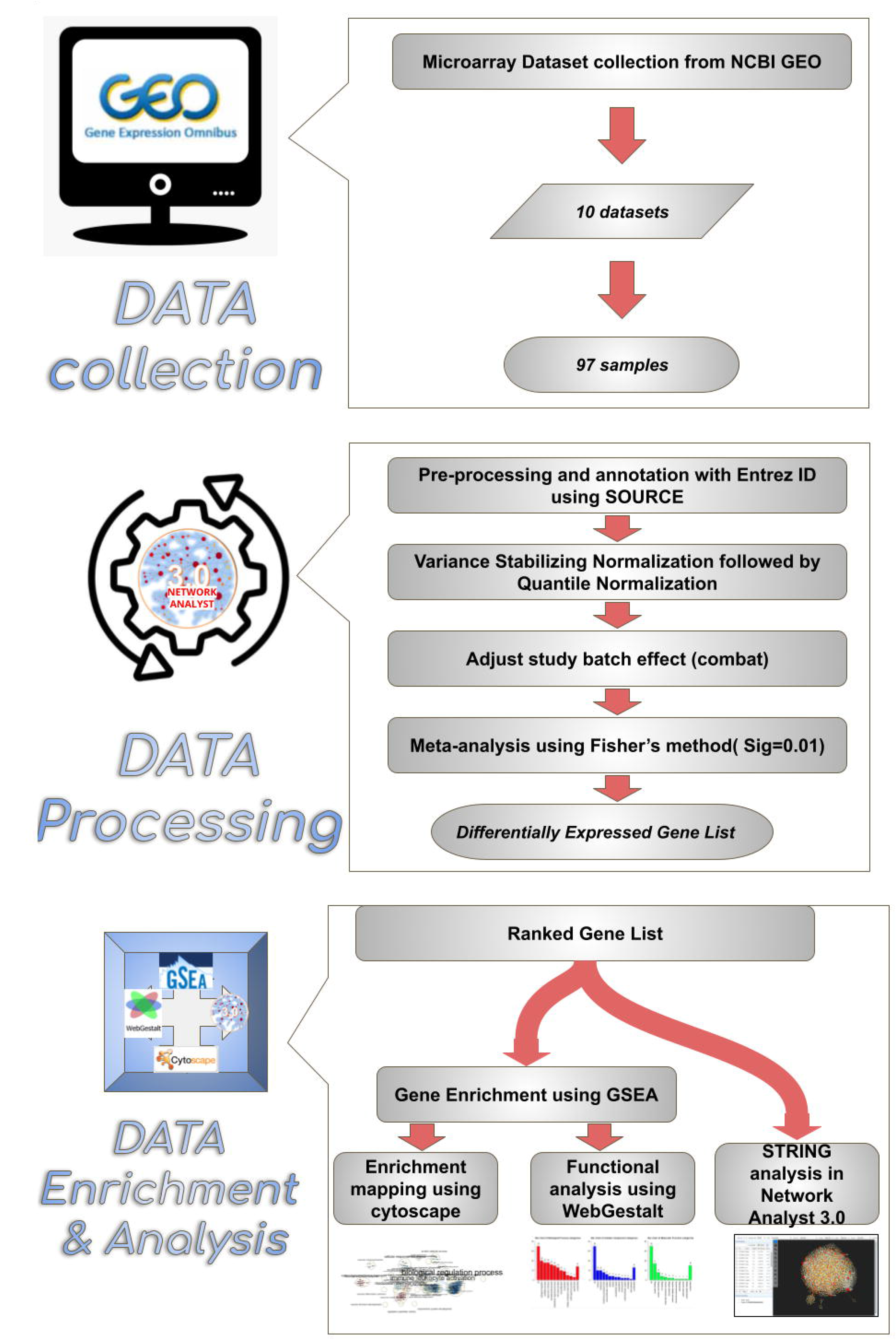
Overview of the enrichment analysis process. Synopsis of meta-analysis conducted in this study. 10 microarray datasets comprising 97 total samples were chosen from NCBI GEO webpage. Datasets were annotated with Entrez id to facilitate compatibility in network analyst 3.0 software. After validation the data were uploaded to Network analyst software, where the samples were normalized and subjected to meta-analysis, thus obtaining the resulting Ranked Differentially expressed gene list comprising 939 genes. The differentially expressed genes between sham and MCAo stroke affected animals were subjected to enrichment analysis through Cytoscape, WebGestalt and string analysis.

**Table-1:** Inclusion Criteria Considered For the Study.

| Inclusion Criteria Considered For the Study |  |
| --- | --- |
| Species | Rattus norvegicus |
| Strain | Sprague-Dawley, Wistar |
| Sex | Male |
| Age | 3 Months - 20 Months (Majority of the sample are in the adult age group ie above 6 months) |
| MCAo Occlusion period | 60 mins, 90 mins, 120 mins, Permanent |
| Sample collection period after MCAo | 3hrs to 7 days (Majority of the sample were collected within the first 3 days) |
| Any model comprising pre-treatment before MCAo was not considered for the study. |  |

**Table-2:** Summary of transcriptome data used in this study.

| Datasets | No.Of Samples |  | Platform | References |
| --- | --- | --- | --- | --- |
|  | Sham | MCAO |  |  |
| GSE97537 | 5 | 7 | GPL1355<br>[Rat230_2] Affymetrix Rat Genome 230 2.0 Array | Unpublished Submission |
| GSE78731 | 5 | 5 | GPL15084<br>Agilent-028279 SurePrint G3 Rat GE 8x60K Microarray (Probe Name version) | (52) |
| GSE52001 | 3 | 3 | GPL14746<br>Agilent-028282 Whole Rat Genome Microarray 4x44K v3 (Probe Name version) | (53) |
| GSE61616 | 5 | 5 | GPL1355<br>[Rat230_2] Affymetrix Rat Genome 230 2.0 Array | (54) |
| GSE33725 | 6 | 6 | GPL7294<br>Agilent-014879 Whole Rat Genome Microarray 4x44K G4131F (Probe Name version) | (55) |
| GSE17929 | 2 | 4 | GPL85 [RG_U34A]<br>Affymetrix Rat Genome U34 Array ///<br>GPL341 [RAE230A]<br>Affymetrix Rat Expression 230A Array | (56) |
| GSE36010 | 8 | 8 | GPL7294<br>Agilent-014879 Whole Rat Genome Microarray 4x44K G4131F (Probe Name version) | (57) |
| GSE23651 | 8 | 8 | GPL6101<br>Illumina ratRef-12 v1.0 expression beadchip | Unpublished Submission |
| GSE46267 | 1 | 8 | GPL6101<br>Illumina ratRef-12 v1.0 expression beadchip | (58) |
| **Total Samples** | 43 | 54 |  |  |

### 3.2) Data processing

Series matrix files were downloaded from the appropriate datasets listed in NCBI and processed by uploading to multiple gene expression table analysis options in Network analyst 3.0. (https://www.networkanalyst.ca/) [4, 5]. For non-compatible datasets, the datasets were processed using SOURCE batch search [6], where the probes from the array chip design were mapped into the appropriate ENTREZ gene ID. The probes that did not match with an Entrez id were not included. Each dataset was individually validated and uploaded to the network analyst software. The study batch effect among different datasets was adjusted using the Combat option [7].

### 3.3) Preparation of differentially expressed genes list

The datasets were normalized by variance stabilizing normalization (VSN) followed by quantile normalization to minimize between slide variations [8]. Then meta-analysis [9] was carried out by combining p-values using Fisher’s method [10] with a significance set at 0.01. The resulting output produced a ranked list of differentially expressed genes. This ranked gene list was further used for pathway enrichment analysis.

### 3.4) Pathway enrichment analysis

The ranked gene list containing upregulated and downregulated genes, and the GO: BP(Gene Ontology: BioProcess) rat GMT file was uploaded to GSEA software for enrichment analysis [11]. The resulting GSEA ranked file was uploaded to the following software modules for various data analyses.

#### 3.4.1) Enrichment map in Cytoscape

To explore the differentially expressed (DE) genes between sham and MCAo groups, the GSEA ranked file was uploaded to the enrichment map application in Cytoscape. Q- value was adjusted to 0.01 and edge cutoff was increased to 0.75. Further, the layout was adjusted to have a defined data and the resulting pathways were annotated. Nodes of interest were selected to observe the list of altered gene lists under the specific gene ontology node. These node-specific gene expression data were also analysed using the Genemania plugin.

#### 3.4.2) WebGestalt

WebGestalt is an online-based enrichment analysis software, useful in analyzing clusters of hits in enriched metabolic and signalling pathways (like KEGG and GO) [12]. Gene set enrichment analysis generated multiple highly significant clusters that showed a common trend as to their function.

#### 3.4.3) Network analyst

Two kinds of analyses were carried out using the DE candidate genes list in the network visual analysis module. Opting for the Transcription Factor (TF)-miRNA interactome analysis reveals non-coding miRNA interaction with the significant genes uploaded. Uploading the data to protein-protein interaction analysis helped in obtaining a string database based interactome map [13].

### 3.5) Venn Diagram Analysis

The differentially expressed gene lists and the highest interacting proteins obtained from the aforementioned analyses were then subjected to venn diagram analysis with previously available gene ontologies related to neurogenesis. The ontology libraries chosen for analyses were i) MANGO (The Mammalian Adult Neurogenesis Gene Ontology), ii) neuroblast proliferation GO:0007405, iii) neuron migration (GO:0001764) iv) neuron differentiation (GO:0030182)

## 4) Results

### 4.1) 939 Differentially expressed genes have been identified in MCAo stroke

An overview of the meta-analysis conducted in this study is shown in Figure-1. The effect of normalization on the datasets had been illustrated in figure-2 exhibiting the difference between pre-normalized and post normalized datasets. Post normalization of the intensities of the achieved datasets was equally distributed amongst the samples well within the median levels. Normalization helps in ensuring minimal experimental variations between the microarray datasets. The normalized datasets were subjected to meta-analysis to identify the differentially expressed genes (DE genes) between MCAo and sham rat brain samples. Fisher’s test with P=0.01 set during the meta-analysis resulted in the identification of 939 DE genes between the groups. The top 50 upregulated, and top 50 downregulated genes in MCAo animals were listed in tables 2 and 3 respectively. Since the current study focuses keenly on identifying the dysregulated genes related to neurogenesis, the DE gene rank list was applied to Gene Set Enrichment Analysis and the enriched gene data was utilized for different visualization and analysis purposes.

**Figure-2:**
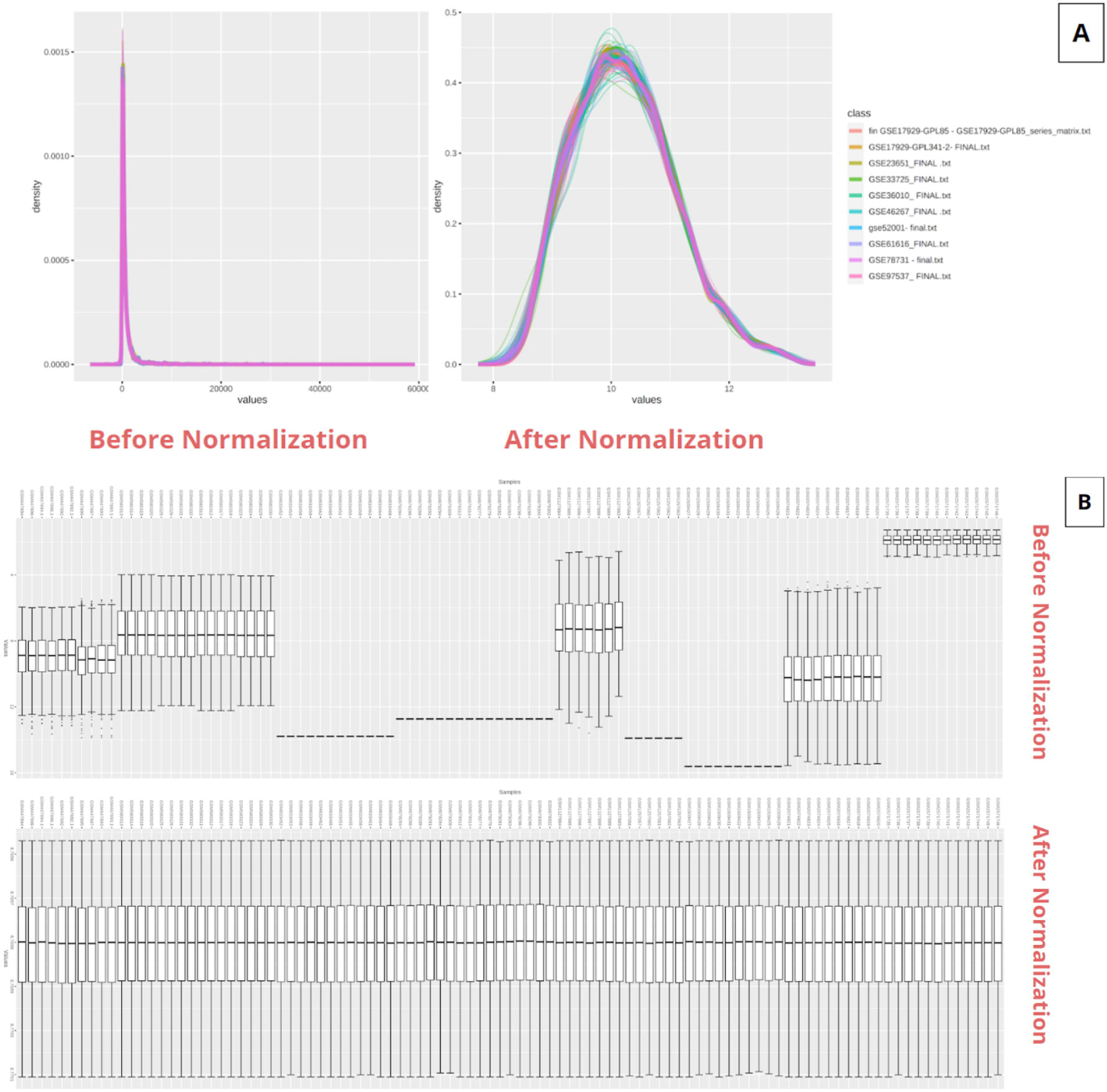
Normalization of Datasets. Normalization of transcriptome data. A) Total intensity distribution among microarray datasets before and after normalization is shown. B) Variation in intensity values from each sample before and after normalization is shown.

**Table-3:** List of top 50 Upregulated genes in stroke brain identified through transcriptome meta-analysis.

| Gene | Name | Entrez id | CombinedTstat |
| --- | --- | --- | --- |
| LGALS3 | galectin 3 | 83781 | -217.89 |
| TSPO | translocator protein | 24230 | -206.96 |
| TIMP1 | TIMP metalloproteinase inhibitor 1 | 116510 | -196.16 |
| CD14 | CD14 molecule [Source:RGD Symbol;Acc:620588] | 60350 | -194.96 |
| HSPB1 | heat shock protein family B (small) member 1 | 24471 | -191.16 |
| GFAP | glial fibrillary acidic protein] | In multiple Geneids | -187.73 |
| CD63 | Cd63 molecule | 29186 | -186.66 |
| C1QC | complement C1q C chain | 362634 | -181.31 |
| HMOX1 | heme oxygenase 1 | 24451 | -180.39 |
| RIPK3 | receptor-interacting serine-threonine kinase 3 | 246240 | -178.3 |
| CCL2 | C-C motif chemokine ligand 2 | 24770 | -172.5 |
| ARPC1B | actin related protein 2/3 complex, subunit 1B | 54227 | -168.72 |
| ATF3 | activating transcription factor 3 | 25389 | -168.66 |
| S100A4 | S100 calcium-binding protein A4 | In multiple Geneids | -168.12 |
| LITAF | lipopolysaccharide-induced TNF factor | In multiple Geneids | -165.64 |
| ANXA1 | annexin A1 | In multiple Geneids | -164.9 |
| HCK | HCK proto-oncogene, Src family tyrosine kinase | 25734 | -164.47 |
| TNFRSF1A | TNF receptor superfamily member 1A | 25625 | -163.26 |
| IFITM3 | interferon induced transmembrane protein 3 | 361673 | -162.77 |
| ICAM1 | intercellular adhesion molecule 1 | 25464 | -160.78 |
| PTPN6 | protein tyrosine phosphatase, non-receptor type 6 | 116689 | -160.7 |
| C1S | complement C1s | 192262 | -159.03 |
| VIM | vimentin | 81818 | -158.62 |
| TBXAS1 | thromboxane A synthase 1 | 24886 | -157.81 |
| GADD45A | growth arrest and DNA-damage-inducible, alpha | 25112 | -157.48 |
| TGFB1 | transforming growth factor, beta 1 | 59086 | -155.89 |
| PSMB9 | proteasome 20S subunit beta 9 | 24967 | -153.44 |
| CAPG | capping actin protein, gelsolin like | 297339 | -153.19 |
| PLA2G4A | phospholipase A2 group IVA | In multiple Geneids | -152.19 |
| IER3 | immediate early response 3 | 294235 | -149.84 |
| CD48 | Cd48 molecule | 245962 | -149.09 |
| GRN | granulin precursor | 29143 | -148.19 |
| CASP1 | caspase 1 | 25166 | -147.52 |
| LY86 | lymphocyte antigen 86 | 291359 | -146.6 |
| CTSL | cathepsin L | 25697 | -146.41 |
| MYC | MYC proto-oncogene, bHLH transcription factor | 24577 | -146.14 |
| STAT3 | signal transducer and activator of transcription 3 | 25125 | -144.69 |
| PSME2 | proteasome activator subunit 2 | 29614 | -144.02 |
| ANXA3 | annexin A3 | 25291 | -140.37 |
| MGP | matrix Gla protein | 25333 | -140.35 |
| PCNA | proliferating cell nuclear antigen | 25737 | -138.9 |
| DYNLT1 | dynein light chain Tctex-type 1 | 83462 | -137.9 |
| IL18 | interleukin 18 | 29197 | -137.5 |
| P2RY6 | pyrimidinergic receptor P2Y6 | 117264 | -136.78 |
| CTSC | cathepsin C | 25423 | -136.49 |
| LGALS3BP | galectin 3 binding protein | 245955 | -132.14 |
| FGR | FGR proto-oncogene, Src family tyrosine kinase | 79113 | -130.06 |
| P4HB | prolyl 4-hydroxylase subunit beta | 25506 | -129.66 |
| RBP1 | retinol binding protein 1 | 25056 | -129.14 |
| CTSB | cathepsin B | 64529 | -128.67 |

**Table-4:** List of top 50 downregulated genes in stroke brain identified through transcriptome meta-analysis.

| Gene | Name | Entrez id | CombinedTstat |
| --- | --- | --- | --- |
| PIK3R2 | phosphoinositide-3-kinase regulatory subunit 2 | 29741 | 127.2 |
| PCSK2 | proprotein convertase subtilisin/kexin type 2 | 25121 | 121.37 |
| B4GALNT1 | beta-1,4-N-acetyl-galactosaminyl transferase 1 | 64828 | 121.04 |
| CX3CL1 | C-X3-C motif chemokine ligand 1 | 89808 | 120.76 |
| SLC6A1 | solute carrier family 6 member 1 | 79212 | 118.59 |
| ATP2B2 | ATPase plasma membrane Ca<sup>2+</sup> transporting 2 | 24215 | 118.25 |
| GRIN1 | glutamate ionotropic receptor NMDA type subunit 1 | 24408 | 117.46 |
| THY1 | Thy-1 cell surface antigen | 24832 | 117.15 |
| KCNV1 | potassium voltage-gated channel modifier subfamily V member 1 | 60326 | 116.7 |
| SCN1B | sodium voltage-gated channel beta subunit 1 | 29686 | 115.96 |
| NEU2 | neuraminidase 2 | 29204 | 115.39 |
| LIN7B | lin-7 homolog B, crumbs cell polarity complex component | 60377 | 114.26 |
| NPY5R | neuropeptide Y receptor Y5 | 25340 | 113.95 |
| ST3GAL2 | ST3 beta-galactoside alpha-2,3-sialyltransferase 2 | 64442 | 113.65 |
| VSNL1 | visinin-like 1 | 24877 | 113.4 |
| ATP1A1 | ATPase Na<sup>+</sup>/K<sup>+</sup> transporting subunit alpha 1 | 24211 | 112.52 |
| CAMK2N1 | calcium/calmodulin-dependent protein kinase II inhibitor 1 | 287005 | 111.85 |
| CLEC11A | C-type lectin domain containing 11A | 29313 | 111.07 |
| PTK2B | protein tyrosine kinase 2 beta | 50646 | 111.03 |
| PDE1B | phosphodiesterase 1B | 29691 | 110.69 |
| DLX5 | distal-less homeobox 5 | 25431 | 110.59 |
| B3GAT1 | beta-1,3-glucuronyltransferase 1 | 117108 | 110.16 |
| PLCB1 | phospholipase C beta 1 | 24654 | 108.99 |
| VAMP1 | vesicle-associated membrane protein 1 | 25624 | 108.88 |
| CHRM4 | cholinergic receptor, muscarinic 4 | 25111 | 108.53 |
| DHCR7 | 7-dehydrocholesterol reductase | 64191 | 107.78 |
| GRIA3 | glutamate ionotropic receptor AMPA type subunit 3 | 29628 | 106.39 |
| SHISA4 | shisa family member 4 | 360848 | 106.14 |
| PDE2A | phosphodiesterase 2A | 81743 | 105.91 |
| PRKAR2B | protein kinase cAMP-dependent type II regulatory subunit beta | 24679 | 105.86 |
| SLC4A3 | solute carrier family 4 member 3 | 24781 | 105.47 |
| NRGN | neurogranin] | 64356 | 105.07 |
| DBN1 | drebrin 1 | 81653 | 104.96 |
| NPTN | neuroplastin | 56064 | 104.93 |
| MCF2L | MCF.2 cell line derived transforming sequence-like | 117020 | 104.78 |
| GRIN2A | glutamate ionotropic receptor NMDA type subunit 2A | 24409 | 103.02 |
| KCNJ4 | potassium inwardly-rectifying channel, subfamily J, member 4 | 116649 | 102.97 |
| KALRN | kalirin, RhoGEF kinase | 84009 | 102.46 |
| SYT2 | synaptotagmin 2 | 24805 | 102.26 |
| GFRA2 | GDNF family receptor alpha 2 | In multiple Geneids | 102.18 |
| OPCML | opioid binding protein/cell adhesion molecule-like | 116597 | 101.75 |
| KCNAB2 | potassium voltage-gated channel subfamily A regulatory beta subunit 2 | 29738 | 101.74 |
| FGF14 | fibroblast growth factor 14 | 63851 | 100.31 |
| NEFM | neurofilament medium | 24588 | 99.746 |
| TRIM54 | tripartite motif-containing 54 | 362708 | 99.65 |
| STXBP1 | syntaxin binding protein 1 | 25558 | 99.618 |
| AKAP5 | A-kinase anchoring protein 5 | 171026 | 98.686 |
| ATP2A2 | ATPase sarcoplasmic/endoplasmic reticulum Ca2+ transporting 2 | 29693 | 98.431 |
| NR4A1 | nuclear receptor subfamily 4, group A, member 1 | 79240 | 97.132 |
| RAB3A | RAB3A, member RAS oncogene family | 25531 | 96.975 |

### 4.2) 35 Neurogenesis related genes have been identified through PPI analysis

Continuation of the analysis module in network analyst through network visual analytics-enabled two types of analysis. One being the transcription factor (TF)- miRNA co-regulatory network analysis (Figure-3), and the second being generic protein-protein interaction analysis (Figure-4). TF-miRNA interactome analysis revealed that TFs like Neurod1, Vimentin, Aquaporin4 were found to be interacting with miRNAs predominantly. These TFs are also crucial during the phases of adult neurogenesis [14–16]. Protein-protein Interaction (PPI) network analysis was also carried out to understand the relation between DE genes based on the intermediate genes of the network. The “zero-order” PPI network contained 163 nodes with 255 interacting edges among them. The most highly interacting node was identified to be Akt1 with betweenness centrality = 5808.71, degree = 22 and Ptk2b With betweenness centrality = 3865.3, degree = 3. A supplementary table with the top 10 interacting nodes is attached (Table-5). In relation to genes related to neurogenesis, around 35 interacting genes were identified by comparing the nodes with GO: Neurogenesis (GO:0022008). The interacting genes related to GO: Neurogenesis were listed in supplementary data (Table-6). Among the proteins related to neurogenesis, the highest interacting among them is found to be Akt1- a known promoter of neurogenesis, Stat3- a known promoter of neuroblast migration, Mtor- a known regulator of neurogenesis, Casp3, Myc- a known promoter of neural differentiation.

**Figure-3:**
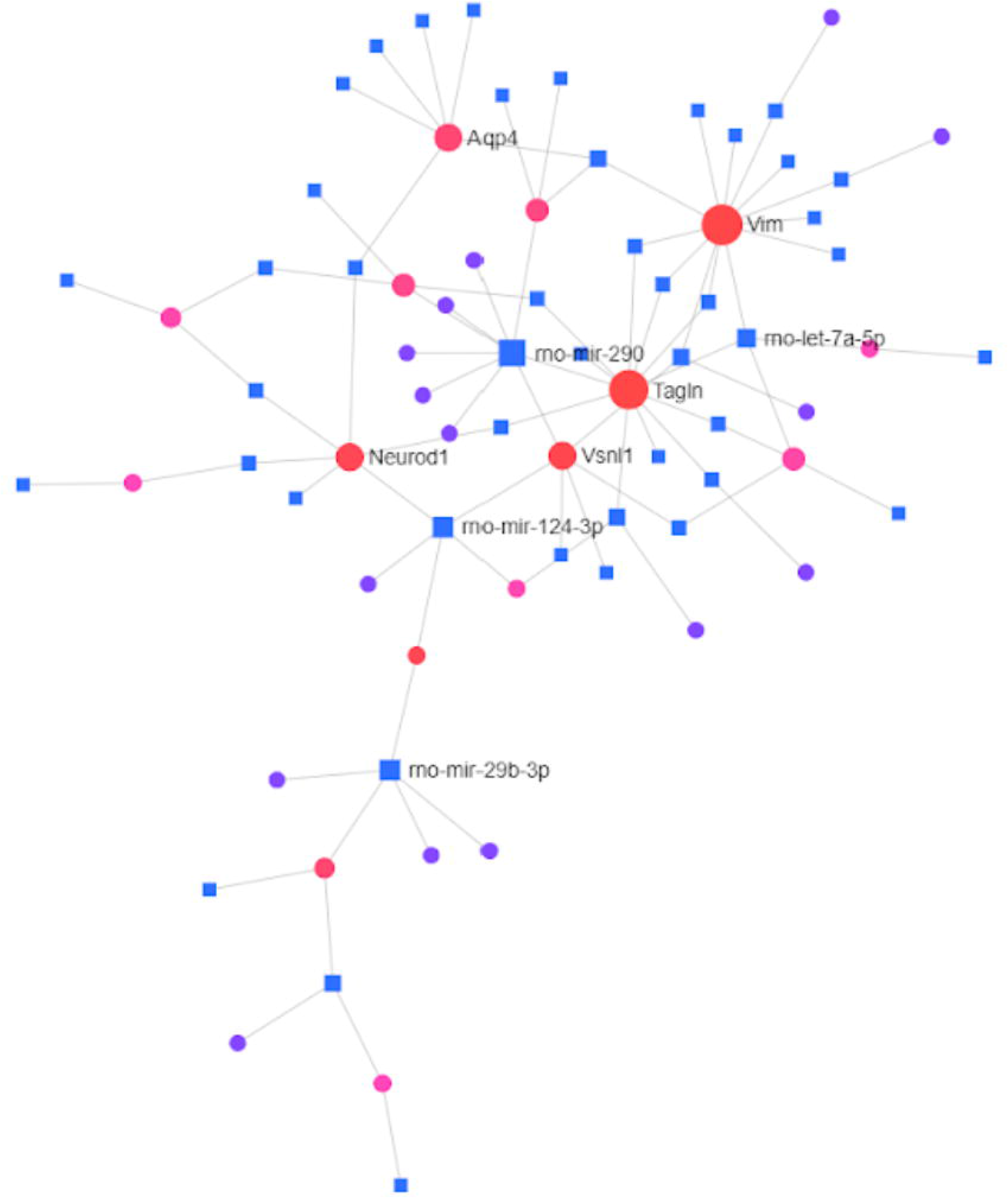
TF- miRNA interactome. TF-miRNA interactome graph. The diagram represents the interaction between miRNA and transcription factors of DE genes where nearest represent those with higher interaction score.

**Figure 4:**
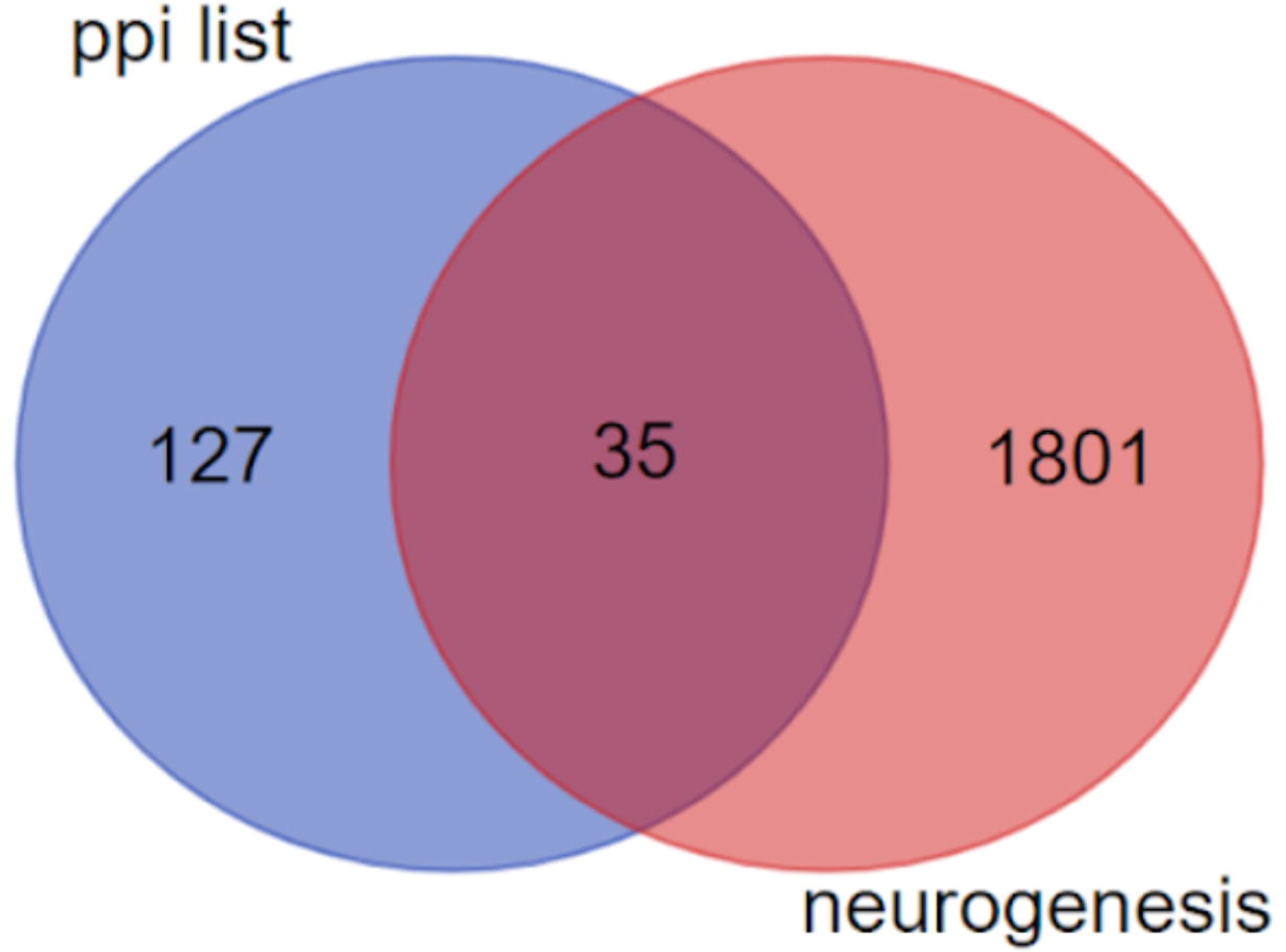
Venn diagram expressing overlapping between PPI list and GO:BP Neurogenesis lists.

**Table-5:** Top 10 Interacting Nodes in PPI Analysis.

| Label | Degree | Betweenness | Expression |
| --- | --- | --- | --- |
| Akt1 | 22 | 5808.71 | 54.118 |
| Cdk1 | 11 | 1780.5 | -106.67 |
| Pcna | 10 | 896.87 | -138.9 |
| Nfkb1 | 9 | 1095.09 | -111.02 |
| Pik3r2 | 8 | 2668.38 | 127.2 |
| Gna11 | 8 | 1679.93 | 92.205 |
| Stat3 | 8 | 1270.92 | -144.69 |
| Syk | 8 | 504.98 | -83.095 |
| Actn2 | 7 | 3294 | 84.253 |
| Hck | 7 | 3016.68 | -164.47 |

**Table-6:**
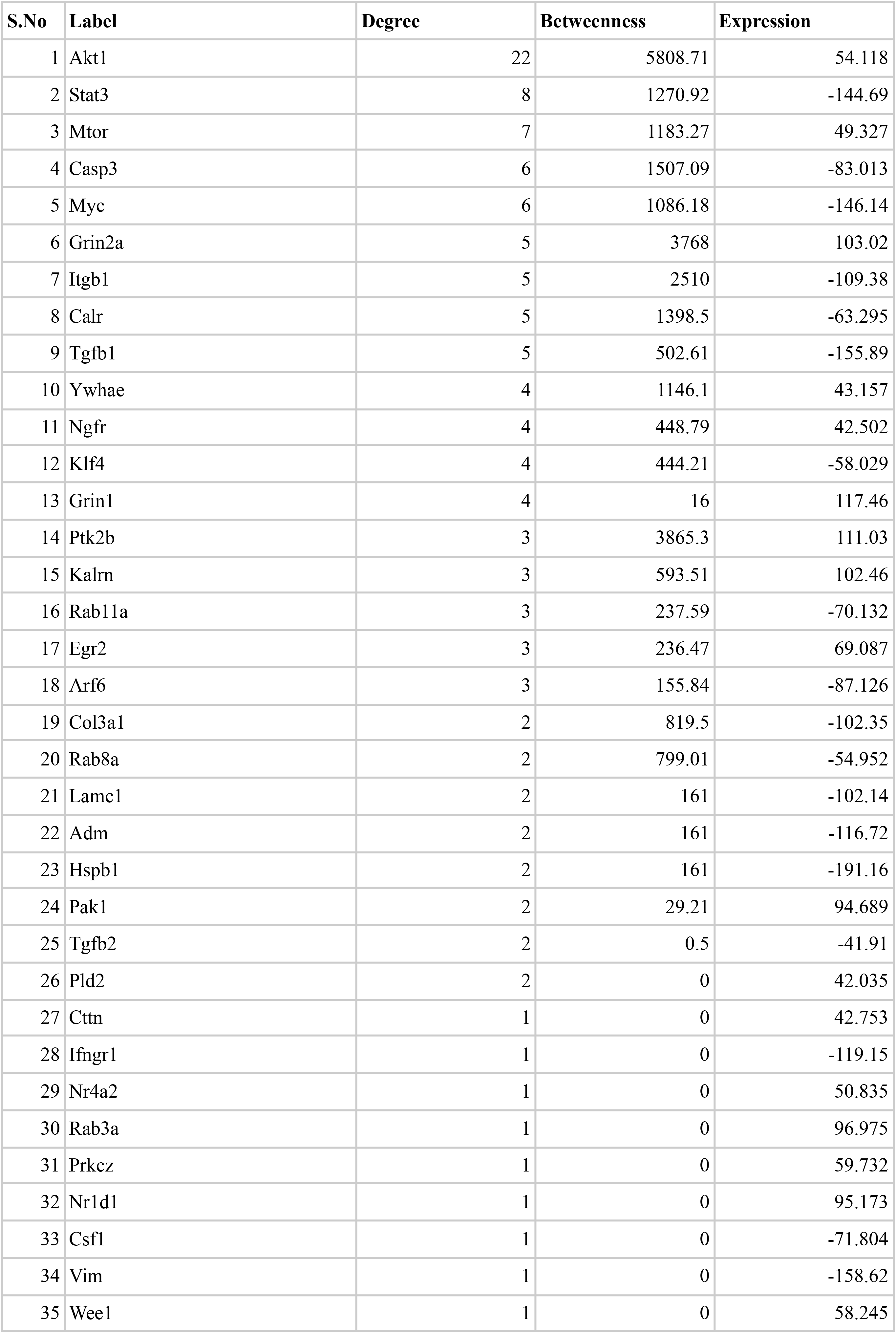
Interacting proteins related to GO: Neurogenesis.

### 4.3) Enrichment analysis using WebGestalt

Moreover, to increase an understanding of the biological roles of DE genes in post MCAo conditions, gene ontology analysis and enrichment analysis were performed in Webgestalt to classify the DE genes into specific readable clusters based on their biological processes, cellular components, and molecular functions during the post MCAo conditions. Gene ontology revealed that biological regulation and localization were predominated with more than 50 % of the DE genes, whereas cell proliferation accounted for less than 15 % of the DE genes. Most of the DE genes were localized to membrane, and nucleus, and fewer in ECM, and ribosomes. Molecular functions related to protein binding were more enriched when compared to oxygen binding and other interactions were least enriched (Fig-5-B). The directed acyclic graph was also constructed to visualize the interaction of enriched gene ontology as schematically depicted in Fig. 5-C. The Kegg based molecular function analysis revealed significant downregulation in nodes like regeneration, immune effector process, while also showing significant upregulation of nodes like regulation of trans-synaptic signalling and glutamate receptor signalling pathway among others (Fig-5-B). Elucidation of the regeneration node in molecular function analysis revealed that neurogenic genes like GFAP, TGFB1, PCNA, JAG1 were significantly upregulated in the MCAo stroke brain.

**Figure-5:**
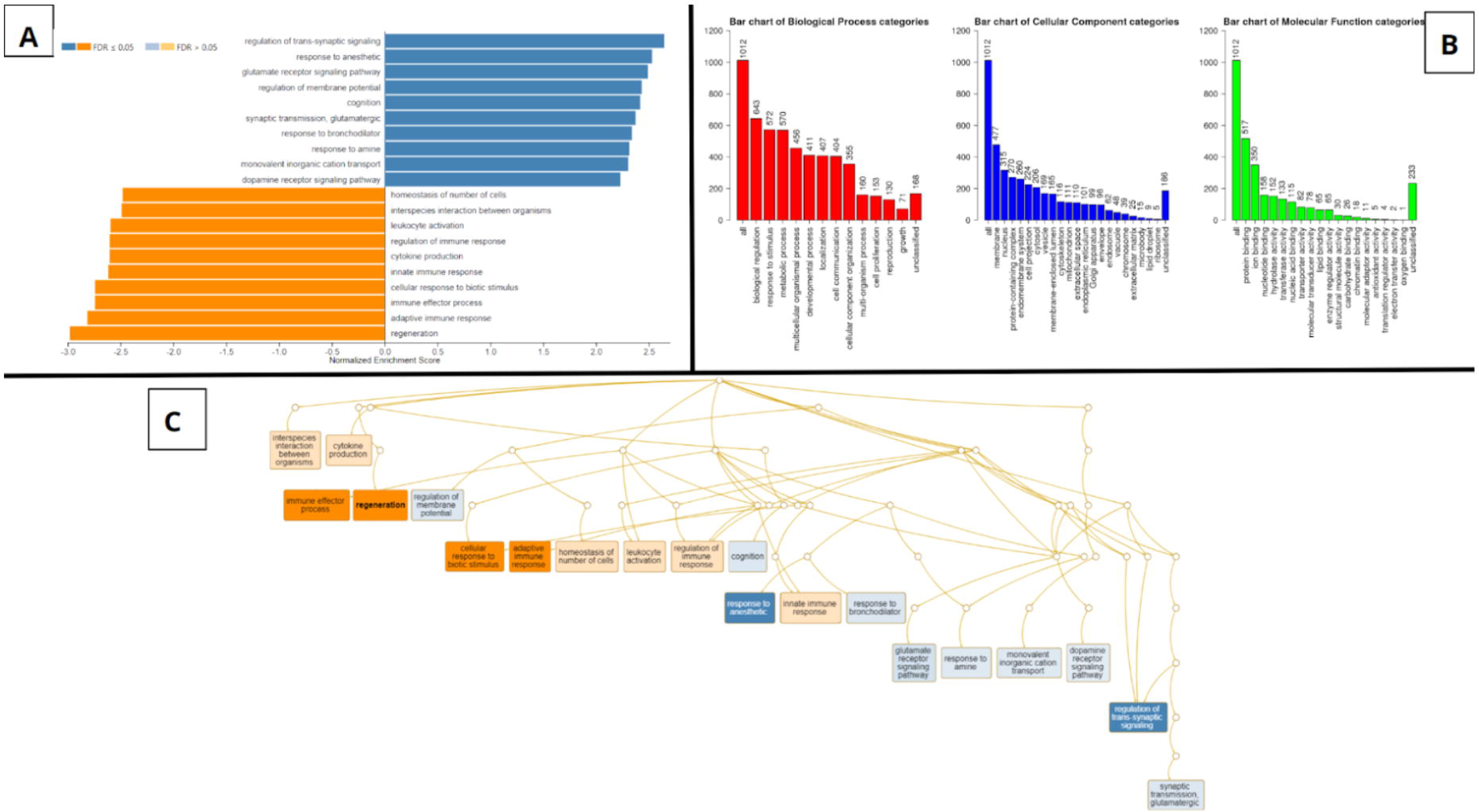
Gene Ontology Classification, Pathway enrichment analysis- Clusters of highly significant pathways based on KEGG Database, Acyclic Graph of enriched processes based on GO Biological Processes. WebGestalt outputs. A) Pathway enrichment analysis of DE genes between MCAo and Sham animals, resulting in multiple clusters of significantly altered pathways based on KEGG database. B) Gene ontology classification. Each bar represents an ontology category and the heigh signifies the number of DE genes observed under each category. C) Branching graph depicts functional interaction between various GO categories, where the steel blue colored categories are positively related while the deep orange colored categories are negatively related.

### 4.4) Enrichment Mapping using Cytoscape

Cytoscape Enrichment map analysis of the GSEA enriched gene list revealed that among various dysregulated pathways, the neurogenesis pathway is one of the significant among them as it passed q value of 0.01 (Fig-6). It depicted that the genes responsible for negative regulation of neurogenesis had been significantly upregulated, which leads to the fact that as whole neurogenic process was impaired in the MCAo group. Genes under the node were displayed in table 7 along with their fold-change score (CombinedTstat), where lower the value corresponds to higher the expression in the MCAo group. The majority of the shortlisted genes have a relation with neurogenesis pathways (Fig-7). Their relevance to different stages of neurogenesis has been discussed ahead.

**Figure-6.**
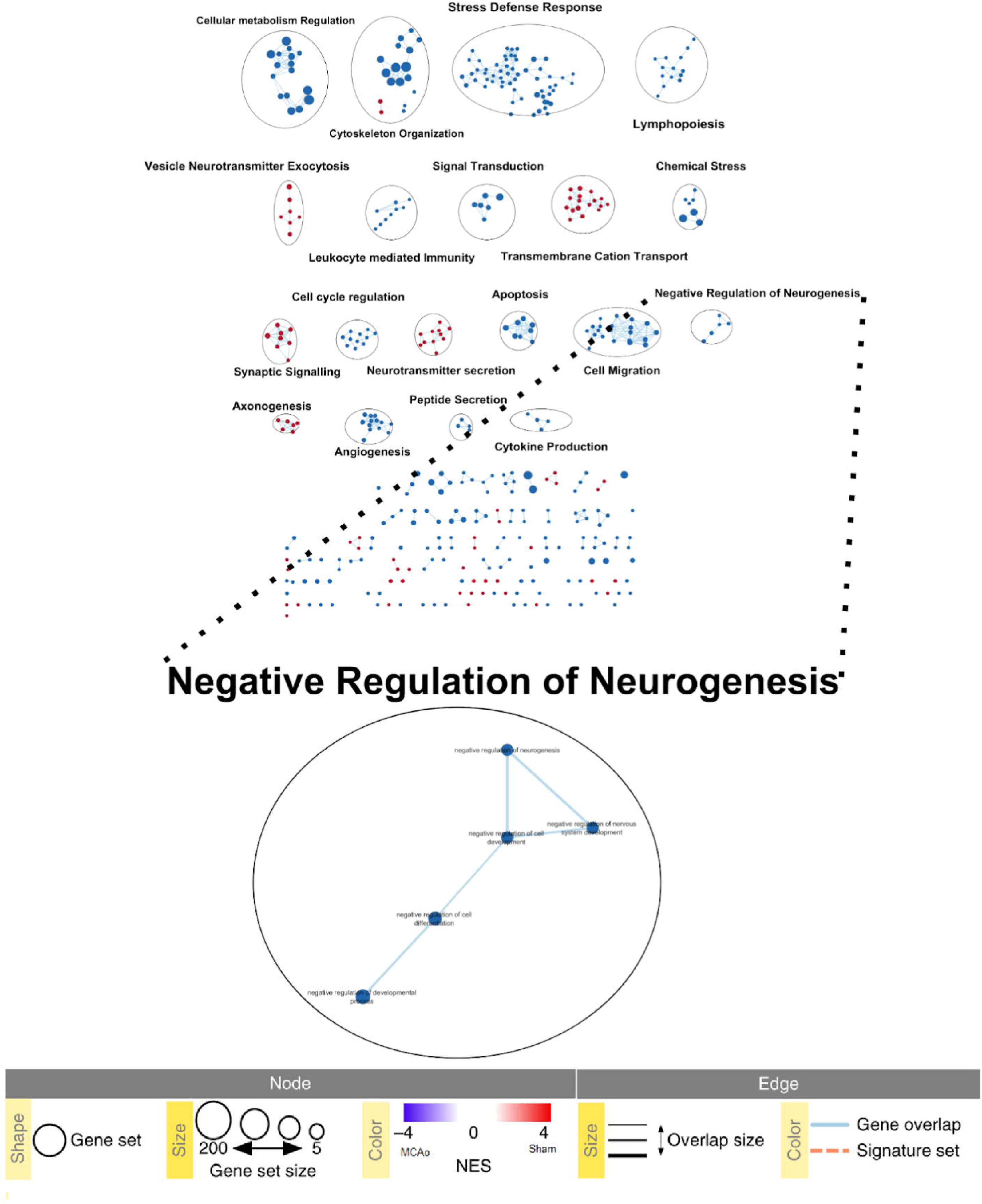
Cytoscape enrichment data. Enrichment map from Cytoscape. The graphic depicts interaction between various enriched nodes(Dots). Lines connecting the nodes are called edge. Blue nodes depict more enrichment in MCAo group while the red nodes depict more enrichment in Sham group.

**Figure-7:**
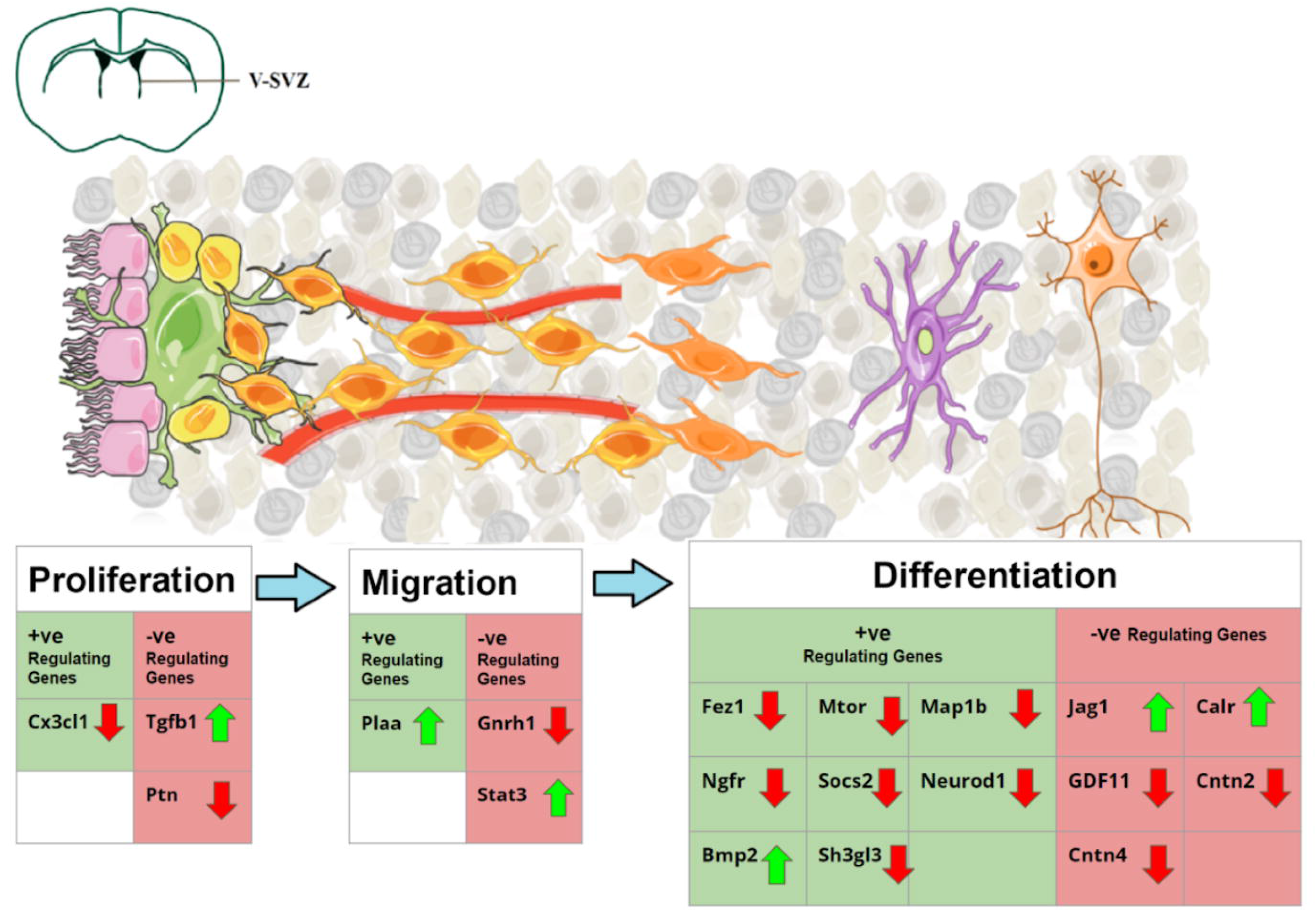
Graphical Representation of the altered neurogenic genes in post stroke environment. Coronal section of the adult rat brain is shown in the upper left. SVZ region comprises Ependymal cells (Pink) lining along the ventricles, with Type B1 stem cells(Green) adhering to them. Type B1 cells give rise to Type-C Transit amplifying Cells(Yellow). During proliferation Type-C cells rapidly divide to form Type-A neuroblasts(Light orange). These neuroblasts migrate along the blood vessels (Red) to reach striatum after stroke. Once reaching the site, the neuroblasts differentiate into immature neurons(violet) and give rise to arborizations. They turn into functional neurons after maturation. The table depicts the genes that upregulate or downregulate the different phases of neurogenesis. The green upward arrows represent genes found to be upregulated in MCAo animals in the meta-analysis study, while the red downward arrow represents genes found to be downregulated in MCAo animals in the meta-analysis study.

**Table-7:**
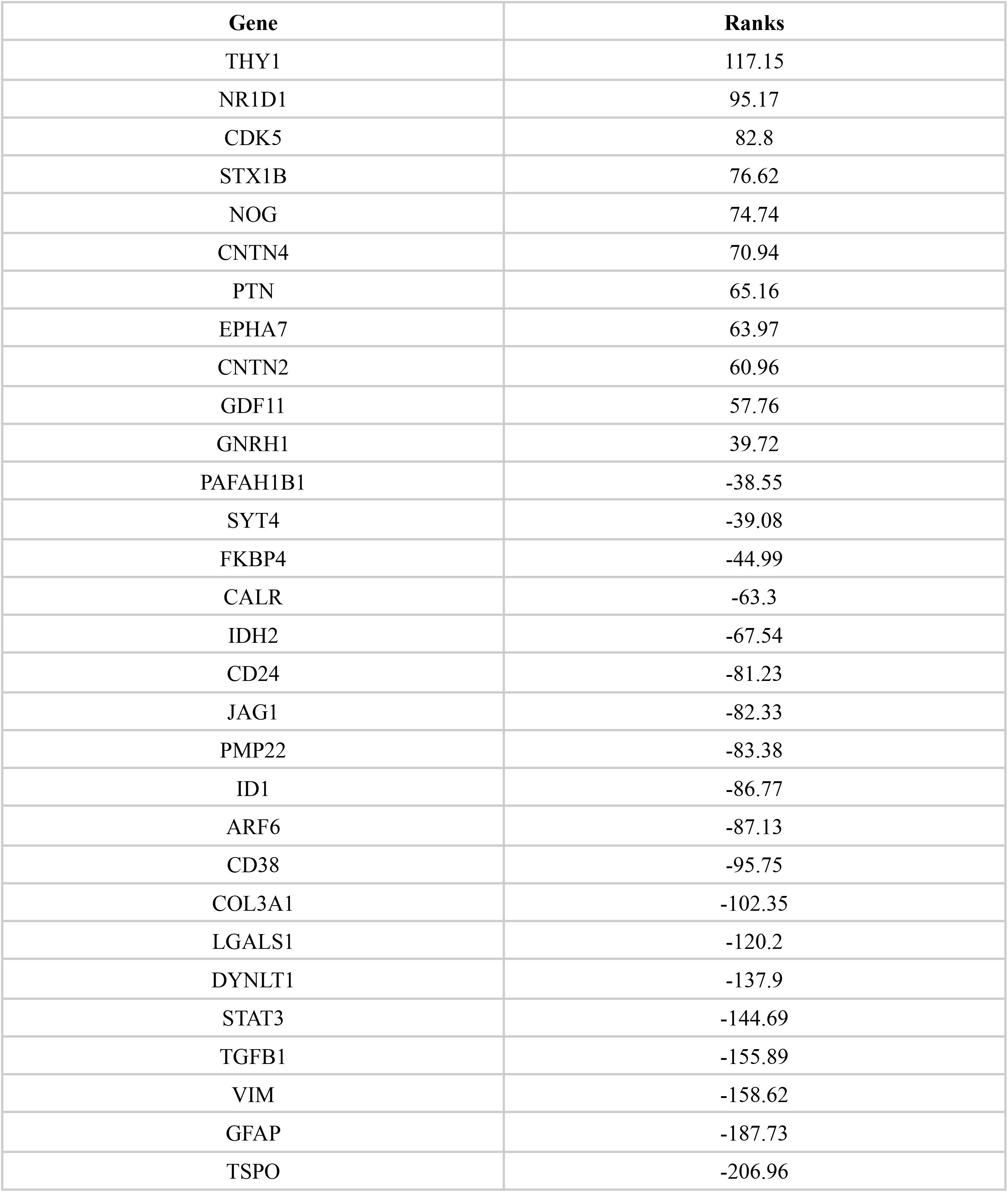
majorly altered genes related to neurogenesis obtained from cytoscape enrichment map.

| **Gene** | **Ranks** |
| --- | --- |
| THY1 | 117.15 |
| NR1D1 | 95.17 |
| CDK5 | 82.8 |
| STX1B | 76.62 |
| NOG | 74.74 |
| CNTN4 | 70.94 |
| PTN | 65.16 |
| EPHA7 | 63.97 |
| CNTN2 | 60.96 |
| GDF11 | 57.76 |
| GNRH1 | 39.72 |
| PAFAH1B1 | -38.55 |
| SYT4 | -39.08 |
| FKBP4 | -44.99 |
| CALR | -63.3 |
| IDH2 | -67.54 |
| CD24 | -81.23 |
| JAG1 | -82.33 |
| PMP22 | -83.38 |
| ID1 | -86.77 |
| ARF6 | -87.13 |
| CD38 | -95.75 |
| COL3A1 | -102.35 |
| LGALS1 | -120.2 |
| DYNLT1 | -137.9 |
| STAT3 | -144.69 |
| TGFB1 | -155.89 |
| VIM | -158.62 |
| GFAP | -187.73 |
| TSPO | -206.96 |

## 5) Discussion

### 5.1) Differentially expressed (DE) Genes in MCAo rat brain when compared with sham

MCAo affected rat brain has an infarct region that encompasses dead cells as a result of the occlusion and the subsequent pathological effects [17]. Several attempts have been made on the development of a treatment strategy to restore lost neurons after stroke. Among the strategies, one of the most capable approaches is the stimulation of neurogenesis to alleviate the neuronal loss sustained in the brain during and after the stroke [18]. Hence in this article, we carry out the identification of dysregulated neurogenic pathways created in post stroke conditions, as this will help in gaining clinically useful insights into a potential therapeutic or diagnostic application. The workflow comprised computation analysis to identify the DE gene list followed by enrichment analysis and subsequent efficient graphical summary for exploratory visualization.

Through a meta-analysis of the datasets, we identified that 939 genes were differentially expressed in the MCAo group in comparison with the sham group. Among them, 369 genes were upregulated and 570 genes were downregulated, both of them significantly with a minimum combined P-value of 5.1732E-6 (Fisher’s 1% test). PIK3R2 was identified to be the highest downregulated gene in this meta-analysis, followed by PCSK2, B4GALNT1, CX3CL1, SLC6A1. The significantly upregulated genes are identified as LGALS3, TSPO, TIMP1, CD14, HSPB1. TF-miRNA interactome analysis revealed that highest interacting transcription factors to be Tagln (Expression: -91.4), Vim (Expression: -158.62), Vsnl1 (Expression: 113.4), Itgb1 (Expression: -109.38), Neurod1 (Expression: 86.53). The highest interacting miRNA was found to be rno-mir-124-3p, rno-mir-290, rno-mir-29b-3p respectively. Through WebGestalt based enrichment analysis, neurogenic genes like GFAP, TGFB1, PCNA, JAG1 were identified to be less enriched in the MCAo rat brain. From these candidates, we could sense a trend that the majority of these genes have a say in the process of neurogenesis as cross-verified using MANGO [19]. In enrichment analysis using Cytoscape, neurogenesis related pathways were chosen. Nodes under negative regulation of neurogenesis were found to be significantly upregulated. Within this node, the differentially altered genes were identified. Significantly downregulated genes were found to be THY1 (Thy-1 Cell Surface Antigen), NR1D1 (Nuclear Receptor Subfamily 1 Group D Member 1), CDK5 (Cyclin-Dependent Kinase 5), STX1B (Syntaxin 1B), NOG (Noggin). Significantly upregulated genes under the neurogenesis node were found to be TSPO (Translocator Protein), GFAP (Glial Fibrillary Acidic Protein), VIM (vimentin), TGFB1 (Transforming Growth Factor Beta 1), STAT3 (Signal Transducer And Activator Of Transcription 3). The results revealed that a significant number of genes and pathways related to neurogenesis were altered during the acute phase after stroke. It is well studied that excitotoxicity caused by stroke leads to a major depletion of neurons in the striatal region [20], but their effect on the neurogenic niche present in the subventricular zone is very feebly understood [21]. This meta-analysis sheds light on the unknown by comparing how the genes and pathways regulating the neurogenic cells in the SVZ region are dysregulated in early post-stroke conditions.

Since the process of SVZ neurogenesis generally comprises the stages- proliferation, migration, differentiation, and maturation, [22] we will individually discuss how the genes regulating each of this phase were found to be altered in this study and the same has been portrayed in the figure 7. To facilitate the discussion, Venn diagram analysis of 939 DE genes with appropriate Gene ontologies as mentioned in methodology was carried out and only the overlapping DE genes had been further discussed here.

### 5.2) SVZ- neuroblast Proliferation

Neuroblast proliferation plays a major role in the production of new neurons as well as the replenishment of the neurogenic niche cells. Any deficiencies to proliferation will lead to a decrease in neuroblast production and subsequent neurogenic niche depletion [23]. These consequences could severely decline the neuroregeneration potential of the brain after stroke. Particularly in this meta-analysis CX3CL1, TGFB1, PTN genes were found to be significantly dysregulated among genes related to neural proliferation. CX3CL1 gene (C-X3-C Motif Chemokine Ligand 1), codes for fractalkine, whose receptors are present in microglia and is crucial for their migration [24]. This gene is significantly downregulated with a combined Tscore of 120.76 in MCAo animals. Being a transmembrane cytokine, CX3CL1 plays a role in communication between the microglia and neuron through its interaction with the CX3CL1 receptor [25]. Microglia plays a crucial role in the survival and proliferation of neural precursor cells [26]. Also, when cx3cl1 is cleaved by α-, β-, and γ-secretase, they create a cx3cl1-ICD which gets translocated into the nucleus to induce survival mode in cells [27]. Fan et al have previously reported that in 5xFAD Tg mice with overexpression of the cx3cl1-ct, a significant reduction in amyloid deposition and reduced neuronal loss, attributed especially to the increased neuroblast proliferation [27]. In post-stroke conditions, decreased cx3cl1 expression could hinder signalling with microglia in addition to disruption of cell survival and neuronal proliferation [28]. TGFB1 (Transforming Growth Factor Beta 1) (CombinedTscore= -155.89), is significantly upregulated in post-stroke conditions. Wachs et al have shown that the addition of TGFb1 to the adult neural stem and progenitor cell cultures (ANPs) resulted in a significant decrease in proliferation i.e. 25% in one-week treatment [29]. Thus TGF-β1 limits the potential of adult neural stem and progenitor cells to proliferate without altering the capacity of self-renewal or the fate of differentiation. PTN (Pleiotrophin) (CombinedTscore= 65.16) gene codes for pleiotrophin, is significantly downregulated in post-stroke conditions. PTN knockout mice (PTN−/−) display increased neuronal stem cell proliferation in the adult mouse cerebral cortex, and exogenous PTN decreases neuronal stem cell proliferation and facilitates cell differentiation [30]. Downregulation of CX3CL1 and upregulation of TGFb1 hinders neural proliferation in post-stroke conditions while downregulation of PTN facilitates a supporting action for neural proliferation.

### 5.3) Neuroblast Migration

After the proliferation of neuroblasts, in the normal brain, they migrate towards the olfactory bulb via the rostral migratory stream (RMS) [22]. However, the previous studies have extensively shown that in conditions like stroke, epilepsy, and other early stages of neurodegenerative diseases, these neuroblasts divert from RMS and invade sites of neuronal damage in and around striatal regions [18]. This happens innately in an attempt to replenish the damages sustained. However, this diverted migration only happens for a short window of time. During the short window of time, the certain proteins that promote neuroblast migration are expressed at the site of damage and beyond this window, their expression is diluted [31]. Any altered expression of genes correlated to neuroblast migration was identified in this meta-analysis, found to be PLAA (Phospholipase A2 Activating Protein), GNRH1(gonadotropin-releasing gonadotropin-releasing hormone 1), and STAT3(signal transducer and activator of transcription 3). PLAA has functions in degradation targeting ubiquitous proteins via the ubiquitin-proteasome network. Due to disrupted post-endocytic trafficking to the lysosome, PLAA KO mice have shown to accumulate dysfunctional VLDLR (Very low-density lipid receptor) in Purkinje cells, which results in disrupted purkinje cell migration as well as their differentiation [32]. In meta-analysis, Plaa(Phospholipase A2 Activating Protein) (CombinedTscore= -74.158), is significantly upregulated in post stroke condition. Higher levels of PLAA expression in post-stroke conditions could lead to improved levels of ubiquitin-protein degradation and could lead to lesser accumulation of disruptive proteins like vldlr which have direct implications in affecting neuroblast migration [33]. To accompany this VLDLR (CombinedTscore= 42.394) expression is downregulated in post-stroke conditions. Larco et al showed that GnRH-(1-5) prevents cellular migration of GN11 cells (GnRH-secreting neuronal cell line) by inhibiting STAT3 signals [34]. STAT3 modulation acts as a mechanism by which extracellular surroundings may control the GnRH neuron migration rate [35]. Meta-analysis revealed significant downregulation of GNRH1 (CombinedTscore= 39.72) and very significant upregulation of STAT3(CombinedTscore= -144.69), both attributing to improvement in the chances of neuroblast migration in the post-stroke brain. In the meta-analysis, significantly altered genes related to neuroblast migration have all cumulatively resulted in promotion of the neuronal migration process. Various other research has shown disruption in slit/robo proteins in post-stroke conditions to be a reason for poor migration after stroke [36]. This meta-analysis provides a new perspective on this matter.

### 5.4) Neuroblast Differentiation and Maturation

Post arrival to the site of damage, it is important for the neuroblast to differentiate into neurons to complement for any functional recovery. Through meta-analysis 13 significantly altered genes related to neuronal differentiation were identified, of which the downregulated genes were NEUROD1(CombinedTscore= 78.801), FEZ1 (CombinedTscore= 78.226), SOCS2(CombinedTscore= 75.286), CNTN4 (CombinedTscore= 58.293), GDF11(CombinedTscore= 57.759), CNTN2 (CombinedTscore= 53.043), MTOR (CombinedTscore= 43.593), SH3GL3(CombinedTscore= 43.274), MAP1B(CombinedTscore= 37.003), and upregulated genes are CALR(CombinedTscore= -71.614), JAG1(CombinedTscore= -68.159), BMP2(CombinedTscore= -57.386), NGFR(CombinedTscore= -49.317). NEUROD1 is one of the well-studied factors in neurogenesis and has been frequently targeted as an option for recovery in neurodegenerative diseases. Gao et al has validated that NEUROD1 ablation in SVZ neurospheres and in-vivo models had resulted in disruption of the survival and differentiation of the newborn neurons [37]. In addition, NeuroD1 specifically binds regulatory elements of neuronal genes, which are silenced by epigenetic pathways in the development, thus proving their importance in neuroblast differentiation [38]. Kang et al had exhibited through shRNA based FEZ1 silencing studies in an in-vivo model that FEZ1 acts synergistically with DISC1 to control the dendritic development of newborn neurons [39]. CNTN2 is a cell-adhesion molecule that is regularly considered as a marker for mature neurons and it is also majorly expressed in regenerating neurons [40]. mTOR signalling is important for neural stem cell maintenance and differentiation and for brain growth [41]. Romine et al have demonstrated that restoration of the mTOR pathway in the ageing brain reestablished neurogenesis [42]. SH3GL3 is a crucial positive regulator of neural differentiation and it interacts with Huntingtin, which has been known to transcriptionally upregulate BDNF expression [43]. Mature neurons consistently express MAP1B and it is a crucial regulator of the structural components of the cell. MAP1B KO resulted in an increased number of excitatory immature symmetrical neurons with altered pre-synaptic modifications [44]. Downregulation of these positive regulators of neural differentiation will severely affect the chances of differentiation in the early phases of post-stroke conditions.

Shih et al showed that the CALR(Calreticulin) deficiency substantially decreased the neuronal differentiation caused by NGF [45], similarly CRT over-expression enhanced neuronal differentiation through simultaneous activation of an ERK-dependent MAPK pathway [46]. However, Dedhar et al have shown that Calreticulin can suppress transcriptional activities of the retinoic acid receptor in vivo, as well as neuronal differentiation caused by retinoic acid [47]. Meta-analyses have revealed that CALR is up-regulated and because of its contrasting roles in the aspect of neural differentiation, a detailed study is needed with respect to SVZ neurogenesis. Nyfeler et al have demonstrated that JAG1 (Jagged1) treatment of NSCs in vitro, substituted for EGF and identified to be a key regulator of NSC- self-renewal and differentiation [48]. However, when astrocytes are co-cultured with neurons, they inhibit neuronal differentiation by endocytosis of Jagged1 through notch pathway [49]. Thus, positive regulation of CALR and JAG1 could have a negative say in the neural differentiation in post-stroke conditions. Yan et al have demonstrated that BMP2 facilitated the differentiation of NSCs into dopaminergic neurons through the involvement of miR-145 and Nurr1 mechanisms [50], thus showing they have a positive say in neuronal differentiation. NGFR (Nerve growth factor receptor) serves as a common receptor for a number of growth factors like BDNF, NGF among others [51] These trophic factors play a crucial role in neural differentiation and the meta-analysis revealing an upregulation of NGFR that would result in an increase in chances of neural differentiation.

## 6) Conclusion

In conclusion, this stringently concorded meta-analysis of differential transcriptomes between sham and MCAo rats helped in the identification of dysregulated genes and pathways in various crucial phases of neurogenesis during early post-stroke conditions. This work acts as a reference for future studies, through which the findings may be validated. Thus, the findings of this study may provide fundamental clues into the pathophysiology of neurogenesis in the early phase after stroke, which could be translated into a therapeutic approach in future.

## Contributions

Conceptualization: SAR, AM; data acquisition: SAR; analysis and interpretation of data: SAR, GE, DG, SKJ, MK, AM; writing–original draft preparation: SAR; writing–review and editing: SAR, SKJ, MK, AM; critical revision of the manuscript for intellectual content: SKJ, MK, AM; supervision: AM

---

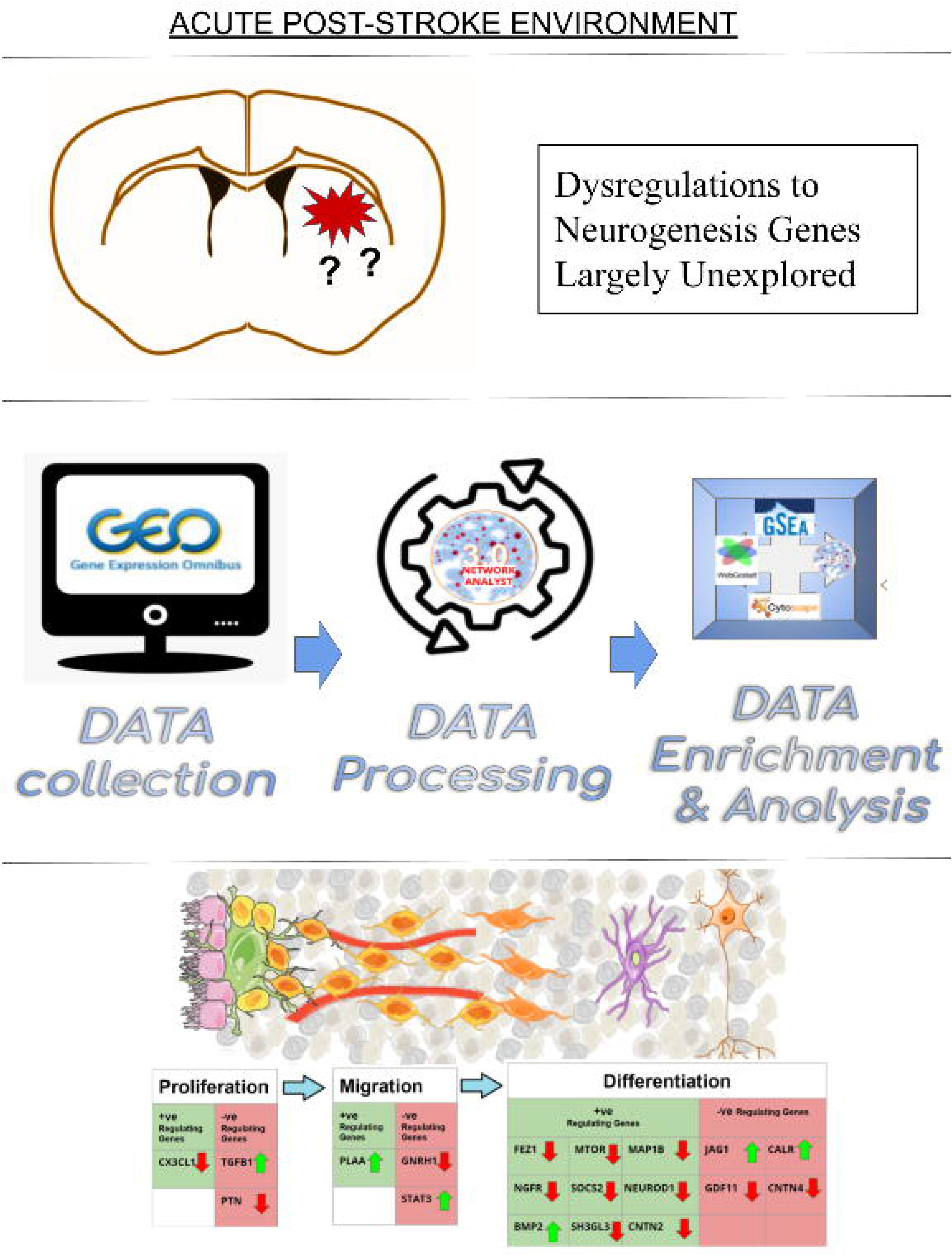

## References

1. Chung, C.-P. (2017). Chapter 77—Types of Stroke and Their Differential Diagnosis. In L. R. Caplan, J. Biller, M. C. Leary, E. H. Lo, A. J. Thomas, M. Yenari, & J. H. Zhang (Eds.), Primer on Cerebrovascular Diseases (Second Edition) (pp. 372–376). San Diego: Academic Press. doi: 10.1016/B978-0-12-803058-5.00077-1

2. Piccardi, B., Arba, F., Nesi, M., Palumbo, V., Nencini, P., Giusti, B., … Inzitari, D. (2018). Reperfusion Injury after ischemic Stroke Study (RISKS): Single-centre (Florence, Italy), prospective observational protocol study. BMJ Open, 8(5). doi: 10.1136/bmjopen-2017-021183

3. Liang, S., Lin, Y., Lin, B., Li, J., Liu, W., Chen, L., … Tao, J. (2017). Resting-state Functional Magnetic Resonance Imaging Analysis of Brain Functional Activity in Rats with Ischemic Stroke Treated by Electro-acupuncture. Journal of Stroke and Cerebrovascular Diseases, 26(9), 1953–1959. doi: 10.1016/j.jstrokecerebrovasdis.2017.06.018

4. Xia, J., Gill, E. E., & Hancock, R. E. W. (2015). NetworkAnalyst for statistical, visual and network-based meta-analysis of gene expression data. Nature Protocols, 10(6), 823–844. doi: 10.1038/nprot.2015.052

5. Zhou, G., Soufan, O., Ewald, J., Hancock, R. E. W., Basu, N., & Xia, J. (2019). NetworkAnalyst 3.0: A visual analytics platform for comprehensive gene expression profiling and meta-analysis. Nucleic Acids Research, 47(W1), W234–W241. doi: 10.1093/nar/gkz240

6. Diehn, M., Sherlock, G., Binkley, G., Jin, H., Matese, J. C., Hernandez-Boussard, T., … Alizadeh, A. A. (2003). SOURCE: A unified genomic resource of functional annotations, ontologies, and gene expression data. Nucleic Acids Research, 31(1), 219–223.

7. Johnson, W. E., Li, C., & Rabinovic, A. (2007). Adjusting batch effects in microarray expression data using empirical Bayes methods. Biostatistics, 8(1), 118–127. doi: 10.1093/biostatistics/kxj037

8. Liu, X., Li, N., Liu, S., Wang, J., Zhang, N., Zheng, X., … Cheng, L. (2019). Normalization Methods for the Analysis of Unbalanced Transcriptome Data: A Review. Frontiers in Bioengineering and Biotechnology, 7. doi: 10.3389/fbioe.2019.00358

9. Xia, J., Fjell, C. D., Mayer, M. L., Pena, O. M., Wishart, D. S., & Hancock, R. E. W. (2013). INMEX—a web-based tool for integrative meta-analysis of expression data. Nucleic Acids Research, 41(W1), W63–W70. doi: 10.1093/nar/gkt338

10. Huo, Z., Tang, S., Park, Y., & Tseng, G. (2020). P-value evaluation, variability index and biomarker categorization for adaptively weighted Fisher’s meta-analysis method in omics applications. Bioinformatics, 36(2), 524–532. doi: 10.1093/bioinformatics/btz589

11. Reimand, J., Isserlin, R., Voisin, V., Kucera, M., Tannus-Lopes, C., Rostamianfar, A., … Bader, G. D. (2019). Pathway enrichment analysis and visualization of omics data using g:Profiler, GSEA, Cytoscape and EnrichmentMap. Nature Protocols, 14(2), 482–517. doi: 10.1038/s41596-018-0103-9

12. Tulalamba, W., Larbcharoensub, N., Sirachainan, E., Tantiwetrueangdet, A., & Janvilisri, T. (2015). Transcriptome meta-analysis reveals dysregulated pathways in nasopharyngeal carcinoma. Tumour Biology: The Journal of the International Society for Oncodevelopmental Biology and Medicine, 36(8), 5931–5942. doi: 10.1007/s13277-015-3268-7.

13. Xia, J., Benner, M. J., & Hancock, R. E. W. (2014). NetworkAnalyst—Integrative approaches for protein–protein interaction network analysis and visual exploration. Nucleic Acids Research, 42(W1), W167–W174. doi: 10.1093/nar/gku443

14. Brulet, R., Zhu, J., Aktar, M., Hsieh, J., & Cho, K.-O. (2017). Mice with conditional NeuroD1 knockout display reduced aberrant hippocampal neurogenesis but no change in epileptic seizures. Experimental Neurology, 293, 190–198. doi: 10.1016/j.expneurol.2017.04.005

15. Morrow, C. S., Porter, T. J., Xu, N., Arndt, Z. P., Ako-Asare, K., Heo, H. J., … Moore, D. L. (2020). Vimentin Coordinates Protein Turnover at the Aggresome during Neural Stem Cell Quiescence Exit. Cell Stem Cell, 26(4), 558–568.e9. doi: 10.1016/j.stem.2020.01.018

16. Liang, R., Yong, S., Huang, X., Kong, H., Hu, G., & Fan, Y. (2016). Aquaporin-4 Mediates the Suppressive Effect of Lipopolysaccharide on Hippocampal Neurogenesis. Neuroimmunomodulation, 23(5–6), 309–317. doi: 10.1159/000467141

17. Fluri, F., Schuhmann, M. K., & Kleinschnitz, C. (2015). Animal models of ischemic stroke and their application in clinical research. Drug Design, Development and Therapy, 9, 3445–3454. doi: 10.2147/DDDT.S56071

18. Lindvall, O., & Kokaia, Z. (2015). Neurogenesis following Stroke Affecting the Adult Brain. Cold Spring Harbor perspectives in biology, 7(11), a019034. https://doi.org/10.1101/cshperspect.a019034

19. Overall RW, Paszkowski-Rogacz M, Kempermann G (2012) The Mammalian Adult Neurogenesis Gene Ontology (MANGO) Provides a Structural Framework for Published Information on Genes Regulating Adult Hippocampal Neurogenesis. PLoS ONE 7(11): e48527. https://doi.org/10.1371/journal.pone.0048527

20. Lima, R. R., Santana, L. N., Fernandes, R. M., Nascimento, E. M., Oliveira, A. C., Fernandes, L. M., Dos Santos, E. M., Tavares, P. A., Dos Santos, I. R., Gimarães-Santos, A., & Gomes-Leal, W. (2016). Neurodegeneration and Glial Response after Acute Striatal Stroke: Histological Basis for Neuroprotective Studies. Oxidative medicine and cellular longevity, 2016, 3173564. https://doi.org/10.1155/2016/3173564

21. Collin, T., Arvidsson, A., Kokaia, Z., & Lindvall, O. (2005). Quantitative analysis of the generation of different striatal neuronal subtypes in the adult brain following excitotoxic injury. Experimental Neurology, 195(1), 71–80. doi: 10.1016/j.expneurol.2005.03.017

22. Alvarez-Buylla, A., & García-Verdugo, J. M. (2002). Neurogenesis in Adult Subventricular Zone. Journal of Neuroscience, 22(3), 629–634. doi: 10.1523/JNEUROSCI.22-03-00629.2002

23. Achanta, P., Capilla-Gonzalez, V., Purger, D., Reyes, J., Sailor, K., Song, H., … Quinones-Hinojosa, A. (2012). Subventricular Zone Localized Irradiation Affects the Generation of Proliferating Neural Precursor Cells and the Migration of Neuroblasts. STEM CELLS, 30(11), 2548–2560. doi: 10.1002/stem.1214

24. Lauro, C., Catalano, M., Trettel, F., Mainiero, F., Ciotti, M. T., Eusebi, F., & Limatola, C. (2006). The chemokine CX3CL1 reduces migration and increases adhesion of neurons with mechanisms dependent on the beta1 integrin subunit. Journal of Immunology (Baltimore, Md.: 1950), 177(11), 7599–7606. doi: 10.4049/jimmunol.177.11.7599

25. Chidambaram, H., Das, R., & Chinnathambi, S. (2020). Interaction of Tau with the chemokine receptor, CX3CR1 and its effect on microglial activation, migration and proliferation. Cell & Bioscience, 10(1), 109. doi: 10.1186/s13578-020-00474-4

26. Chounchay, S., Noctor, S. C., & Chutabhakdikul, N. (2020). Microglia enhances proliferation of neural progenitor cells in an in vitro model of hypoxic-ischemic injury. EXCLI Journal, 19, 950–961. doi: 10.17179/excli2020-2249

27. Fan, Q., Gayen, M., Singh, N., Gao, F., He, W., Hu, X., … Yan, R. (2019a). The intracellular domain of CX3CL1 regulates adult neurogenesis and Alzheimer’s amyloid pathology. The Journal of Experimental Medicine, 216(8), 1891–1903. doi: 10.1084/jem.20182238.

28. He, H.-Y., Ren, L., Guo, T., & Deng, Y.-H. (2019). Neuronal autophagy aggravates microglial inflammatory injury by downregulating CX3CL1/fractalkine after ischemic stroke. Neural Regeneration Research, 14(2), 280–288. doi: 10.4103/1673-5374.244793

29. Wachs, F.-P., Winner, B., Couillard-Despres, S., Schiller, T., Aigner, R., Winkler, J., … Aigner, L. (2006). Transforming Growth Factor-β1 Is a Negative Modulator of Adult Neurogenesis. Journal of Neuropathology & Experimental Neurology, 65(4), 358–370. doi: 10.1097/01.jnen.0000218444.53405.f0

30. Tang, C., Wang, M., Wang, P., Wang, L., Wu, Q., & Guo, W. (2019). Neural Stem Cells Behave as a Functional Niche for the Maturation of Newborn Neurons through the Secretion of PTN. Neuron, 101(1), 32–44.e6. doi: 10.1016/j.neuron.2018.10.051

31. Thored Pär, Wood James, Arvidsson Andreas, Cammenga Jörg, Kokaia Zaal, & Lindvall Olle. (2007). Long-Term Neuroblast Migration Along Blood Vessels in an Area With Transient Angiogenesis and Increased Vascularization After Stroke. Stroke, 38(11), 3032–3039. doi: 10.1161/STROKEAHA.107.488445

32. Hall, E. A., Nahorski, M. S., Murray, L. M., Shaheen, R., Perkins, E., Dissanayake, K. N., Kristaryanto, Y., Jones, R. A., Vogt, J., Rivagorda, M., Handley, M. T., Mali, G. R., Quidwai, T., Soares, D. C., Keighren, M. A., McKie, L., Mort, R. L., Gammoh, N., Garcia-Munoz, A., Davey, T., … Mill, P. (2017). PLAA Mutations Cause a Lethal Infantile Epileptic Encephalopathy by Disrupting Ubiquitin-Mediated Endolysosomal Degradation of Synaptic Proteins. American journal of human genetics, 100(5), 706–724. https://doi.org/10.1016/j.ajhg.2017.03.008

33. DiBattista, A. M., Dumanis, S. B., Song, J. M., Bu, G., Weeber, E., Rebeck, G. W., & Hoe, H. S. (2015). Very low density lipoprotein receptor regulates dendritic spine formation in a RasGRF1/CaMKII dependent manner. Biochimica et biophysica acta, 1853(5), 904–917. https://doi.org/10.1016/j.bbamcr.2015.01.015

34. Larco, D. O., Semsarzadeh, N. N., Cho-Clark, M., Mani, S. K., & John Wu, T. (2013). The Novel Actions of the Metabolite GnRH-(1-5) are Mediated by a G Protein-Coupled Receptor. Frontiers in Endocrinology, 4. doi: 10.3389/fendo.2013.00083

35. Wierman, M. E., Kiseljak-Vassiliades, K., & Tobet, S. (2011). Gonadotropin Releasing Hormone (GnRH) Neuron Migration: Initiation, Maintenance and Cessation as Critical Steps to Ensure Normal Reproductive Function. Frontiers in Neuroendocrinology, 32(1), 43–52. doi: 10.1016/j.yfrne.2010.07.005

36. Naoko, K., Herranz-Pérez, V., Otsuka, T., Sano, H., Ohno, N., Omata, T., … Sawamoto, K. (2018). New neurons use Slit-Robo signaling to migrate through the glial meshwork and approach a lesion for functional regeneration. Science Advances, 4, eaav0618. doi: 10.1126/sciadv.aav0618

37. Gao, Z., Ure, K., Ables, J. L., Lagace, D. C., Nave, K.-A., Goebbels, S., … Hsieh, J. (2009). Neurod1 is essential for the survival and maturation of adult-born neurons. Nature Neuroscience, 12(9), 1090–1092. doi: 10.1038/nn.2385

38. Pataskar, A., Jung, J., Smialowski, P., Noack, F., Calegari, F., Straub, T., & Tiwari, V. K. (2016). NeuroD1 reprograms chromatin and transcription factor landscapes to induce the neuronal program. The EMBO Journal, 35(1), 24–45. doi: 10.15252/embj.201591206

39. Kang, E., Burdick, K. E., Kim, J. Y., Duan, X., Guo, J. U., Sailor, K. A., … Ming, G. (2011). Interaction between FEZ1 and DISC1 in regulation of neuronal development and risk for schizophrenia. Neuron, 72(4), 559–571. doi: 10.1016/j.neuron.2011.09.032

40. Kastriti, M. E., Stratigi, A., Mariatos, D., Theodosiou, M., Savvaki, M., Kavkova, M., … Karagogeos, D. (2019). Ablation of CNTN2+ Pyramidal Neurons During Development Results in Defects in Neocortical Size and Axonal Tract Formation. Frontiers in Cellular Neuroscience, 13. doi: 10.3389/fncel.2019.00454

41. Lee, D. Y. (2015). Roles of mTOR Signaling in Brain Development. Experimental Neurobiology, 24(3), 177–185. doi: 10.5607/en.2015.24.3.177

42. Romine, J., Gao, X., Xu, X.-M., So, K. F., & Chen, J. (2015). The proliferation of amplifying neural progenitor cells is impaired in the aging brain and restored by the mTOR pathway activation. Neurobiology of Aging, 36(4), 1716–1726. doi: 10.1016/j.neurobiolaging.2015.01.003

43. Sari Y. (2011). Huntington’s Disease: From Mutant Huntingtin Protein to Neurotrophic Factor Therapy. International journal of biomedical science : IJBS, 7(2), 89–100.

44. Bodaleo, F. J., Montenegro-Venegas, C., Henríquez, D. R., Court, F. A., & Gonzalez-Billault, C. (2016). Microtubule-associated protein 1B (MAP1B)-deficient neurons show structural presynaptic deficiencies in vitro and altered presynaptic physiology. Scientific Reports, 6. doi: 10.1038/srep30069

45. Lee, A. C.-L., Shih, Y.-Y., Zhou, F., Chao, T.-C., Lee, H., Liao, Y.-F., … Hong, J.-H. (2019). Calreticulin regulates MYCN expression to control neuronal differentiation and stemness of neuroblastoma. Journal of Molecular Medicine, 97(3), 325–339. doi: 10.1007/s00109-018-1730-x

46. Shih, Y.-Y., Nakagawara, A., Lee, H., Juan, H.-F., Jeng, Y.-M., Lin, D.-T., … Liao, Y.-F. (2012). Calreticulin mediates nerve growth factor-induced neuronal differentiation. Journal of Molecular Neuroscience: MN, 47(3), 571–581. doi: 10.1007/s12031-011-9683-3

47. Dedhar, S., Rennie, P. S., Shago, M., Hagesteijn, C.-Y. L., Yang, H., Filmus, J., … Giguère, V. (1994). Inhibition of nuclear hormone receptor activity by calreticulin. Nature, 367(6462), 480–483. doi: 10.1038/367480a0.

48. Wilhelmsson, U., Faiz, M., Pablo, Y. de, Sjöqvist, M., Andersson, D., Widestrand, Å., … Pekny, M. (2012). Astrocytes Negatively Regulate Neurogenesis Through the Jagged1-Mediated Notch Pathway. STEM CELLS, 30(10), 2320–2329. doi: 10.1002/stem.1196

49. Yan, W., Chen, Z.-Y., Chen, J.-Q., & Chen, H.-M. (2016). BMP2 promotes the differentiation of neural stem cells into dopaminergic neurons in vitro via miR-145-mediated upregulation of Nurr1 expression. American Journal of Translational Research, 8(9), 3689–3699.

50. Tomellini, E., Lagadec, C., Polakowska, R., & Le Bourhis, X. (2014). Role of p75 neurotrophin receptor in stem cell biology: More than just a marker. Cellular and Molecular Life Sciences, 71(13), 2467–2481. doi: 10.1007/s00018-014-1564-9

51. Oh, S.-H., Choi, C., Noh, J.-E., Lee, N., Jeong, Y.-W., Jeon, I., … Song, J. (2018). Interleukin-1 receptor antagonist-mediated neuroprotection by umbilical cord-derived mesenchymal stromal cells following transplantation into a rodent stroke model. Experimental & Molecular Medicine, 50(4), 22. doi: 10.1038/s12276-018-0041-1

52. Lai, W., Zheng, Z., Zhang, X., Wei, Y., Chu, K., Brown, J., … Chen, L. (2015). Salidroside-Mediated Neuroprotection is Associated with Induction of Early Growth Response Genes (Egrs) Across a Wide Therapeutic Window. Neurotoxicity Research, 28(2), 108–121. doi: 10.1007/s12640-015-9529-9

53. Wang, L., Yu, Y., Yang, J., Zhao, X., & Li, Z. (2015). Dissecting Xuesaitong’s mechanisms on preventing stroke based on the microarray and connectivity map. Molecular BioSystems, 11(11), 3033–3039. doi: 10.1039/c5mb00379b

54. Mengozzi, M., Cervellini, I., Villa, P., Erbayraktar, Z., Gökmen, N., Yilmaz, O., … Ghezzi, P. (2012). Erythropoietin-induced changes in brain gene expression reveal induction of synaptic plasticity genes in experimental stroke. Proceedings of the National Academy of Sciences of the United States of America, 109(24), 9617–9622. doi: 10.1073/pnas.1200554109

55. Armugam, A., Cher, C. D. N., Lim, K., Koh, D. C. I., Howells, D. W., & Jeyaseelan, K. (2009). A secretory phospholipase A2-mediated neuroprotection and anti-apoptosis. BMC Neuroscience, 10, 120. doi: 10.1186/1471-2202-10-120

56. Ramos-Cejudo, J., Gutiérrez-Fernández, M., Rodríguez-Frutos, B., Expósito Alcaide, M., Sánchez-Cabo, F., Dopazo, A., & Díez-Tejedor, E. (2012). Spatial and temporal gene expression differences in core and periinfarct areas in experimental stroke: A microarray analysis. PloS One, 7(12), e52121. doi: 10.1371/journal.pone.0052121

57. Liu, F. J., Lim, K. Y., Kaur, P., Sepramaniam, S., Armugam, A., Wong, P. T. H., & Jeyaseelan, K. (2013). MicroRNAs Involved in Regulating Spontaneous Recovery in Embolic Stroke Model. PloS One, 8(6), e66393. doi: 10.1371/journal.pone.0066393

